# Negligible-Cost and Weekend-Free Chemically Defined Human iPSC Culture

**DOI:** 10.1101/685503

**Authors:** Hui-Hsuan Kuo, Xiaozhi Gao, Jean-Marc DeKeyser, K. Ashley Fetterman, Emily A. Pinheiro, Carly J. Weddle, Hananeh Fonoudi, Michael V. Orman, Marisol Romero-Tejeda, Mariam Jouni, Malorie Blancard, Tarek Magdy, Conrad L. Epting, Alfred L. George, Paul W. Burridge

## Abstract

Human induced pluripotent stem cell (hiPSC) culture has become routine, yet pluripotent cell media costs, frequent media changes, and reproducibility of differentiation have remained restrictive, limiting the potential for large-scale projects. Here, we describe the formulation of a novel hiPSC culture medium (B8) as a result of the exhaustive optimization of medium constituents and concentrations, establishing the necessity and relative contributions of each component to the pluripotent state and cell proliferation. B8 eliminates 97% of the costs of commercial media, made possible primarily by the in-lab generation of three *E. coli*-expressed, codon-optimized recombinant proteins: an engineered form of fibroblast growth factor 2 (FGF2) with improved thermostability (FGF2-G3); transforming growth factor β3 (TGFβ3) - a more potent TGFβ able to be expressed in *E. coli*; and a derivative of neuregulin 1 (NRG1) containing the EGF-like domain. The B8 formula is specifically optimized for fast growth and robustness at low seeding densities. We demonstrated the derivation and culture of 34 hiPSC lines in B8 as well as maintenance of pluripotency long-term (over 100 passages). This formula also allows a weekend-free feeding schedule without sacrificing growth rate or capacity for differentiation. Thus, this simple, cost-effective, and open source B8 media, will enable large hiPSC disease modeling projects such as those being performed in pharmacogenomics and large-scale cell production required for regenerative medicine.

## Introduction

Human induced pluripotent stem cells (hiPSCs) are functionally immortal and can proliferate without limit while maintaining the potential to differentiate to, hypothetically, all ~220 cell lineages within the human body. hiPSC generation has become routine due to the simplicity of amplification of CD71^+^ blood proerythroblasts (Chou et al., 2015; Chou et al., 2011; Tan et al., 2014) or myeloid cells (Eminli et al., 2009; Staerk et al., 2010) and commercial Sendai virus-based reprogramming factor expression (Fujie et al., 2014; Fusaki et al., 2009). This simplicity has resulted in increased enthusiasm for the potential applications of hiPSC-derived cells across many fields, including regenerative medicine, disease modeling, drug discovery, and pharmacogenomics.

However, these applications require the culture of either large quantities of hiPSCs or hiPSC lines derived from large numbers of patients, and three major restrictions have become evident: 1, the cost of large-scale pluripotent cell culture, which is prohibitive for high patient-number projects; 2, the time-consuming requirement for daily media changes, which is particularly problematic for laboratories in industry; 3, inter-line variability in differentiation efficacy, which is highly dependent on pluripotent culture consistency and methodology. Our understanding of the media conditions required to culture human pluripotent stem cells has progressed steadily over the last 15 years, with significant breakthroughs coming from the discovery of the necessity for high concentrations of FGF2 (Xu et al., 2005), the use of TGFβ1 (Amit et al., 2004), and the elimination of knockout serum replacement (KSR) with the TeSR formula which contains 19 components (Ludwig and Thomson, 2007; Ludwig et al., 2006a; Ludwig et al., 2006b), followed by the first robust chemically defined formula, E8 (Beers et al., 2012; Chen et al., 2011) which consists of just 8 major components. A number of alternative non-chemically defined pluripotent formulations have been described including: CDM-BSA (Hannan et al., 2013; Vallier et al., 2005; Vallier et al., 2009), DC-HAIF (Singh et al., 2012; Wang et al., 2007), hESF9T (Furue et al., 2008; Yamasaki et al., 2014), FTDA (Breckwoldt et al., 2017; Frank et al., 2012; Piccini et al., 2015), and iDEAL (Marinho et al., 2015) (Figure S1A).

Each of the available formulations consist of a core of three major signaling components: 1, insulin or IGF1 which bind INSR and IGF1R to signal the PI3K/AKT pathway promoting survival and growth; 2, FGF2 and/or NRG1 which bind FGFR1/FGFR4 or ERBB3/ERBB4 respectively, activating the PI3K/AKT/mTOR and MAPK/ERK pathways; and 3, TGFβ1, NODAL, or activin A which bind TGFBR1/2 and/or ACVR2A/2B/1B/1C to activate the TGFβ signaling pathway. NODAL is used less commonly in pluripotent media formulations due to the expression of NODAL antagonists LEFTY1/2 in human pluripotent stem cells (hPSC) (Besser, 2004; Sato et al., 2003) resulting in a requirement for high concentrations *in vitro* (Chen et al., 2011). In addition, numerous growth factor-free formulae utilizing small molecules to replace some or all growth factors in hPSC culture have been described (Burton et al., 2010; Desbordes et al., 2008; Kumagai et al., 2013; Tsutsui et al., 2011), however, these have not successfully translated to common usage. Recently, a growth factor-free formula AKIT was demonstrated (Yasuda et al., 2018), combining inhibitors of GSK3B (1-azakenpaulone), DYRK1 (ID-8), and calcineurin/NFAT (tacrolimus/FK506), albeit with much reduced proliferation and colony growth, as well as increased interline variability in growth. Finally, more than 15 commercial pluripotent media are also available in which the formulae are proprietary and not disclosed to researchers. These media represent the major cost for most hiPSC labs and considerably restrict research efforts. Some of these media formulae are suggested to support hiPSC growth without daily media changes or ‘weekend-free’, likely by using heparin sulfate to stabilize FGF2 that otherwise degrades quickly at 37 °C (Chen et al., 2012; Furue et al., 2008) and including bovine serum albumin (BSA) which acts as a multifaceted antioxidant.

Here, we demonstrate a novel media formula (B8), thoroughly optimized to support high growth rate under low seeding density conditions, that requires minimal media exchanges, is low cost, and maintains differentiation reproducibility. This formula is capable of supporting both hiPSC generation and long-term culture for >100 passages. Production of B8 supplement aliquots suitable for making 100 liters of B8 medium is simple for any research lab with basic equipment, with complete bottles of medium costing ~$16 USD per liter. A full protocol is provided, including detailed instructions for recombinant protein production in three simple steps. All plasmids for protein production are available through Addgene. With the commoditization of these protocols, we believe it is possible to substantially increase what is achievable with hiPSCs due to the near elimination of pluripotent cell culture costs and minimization of labor associated with cell culture.

## Results

### Optimization of Media Constituents with a Short-Term Assay

We began with simple cost-reductions to our existing hiPSC culture medium formula based on E8 that we have used extensively (Figure 1A). First, we optimized suitable concentrations of matrices on which the hiPSCs are grown. Although laminin-511 (Rodin et al., 2010), laminin-521 (Rodin et al., 2014), vitronectin (Braam et al., 2008), and Synthemax-II (Melkoumian et al., 2010) are suitable for hiPSC culture (Burridge et al., 2014), none are appropriately cost-effective, and in the case of vitronectin or Synthemax-II, also not suitable for subsequent cardiomyocyte differentiation (Burridge et al., 2014). Matrigel, although an undefined product (Hughes et al., 2010), is a cost-effective and commonly used matrix with substantial data using it at 50 µg cm^−2^ (Ludwig et al., 2006a). Comparing two similar commercial products, Matrigel (Corning) and Cultrex/Geltrex (Trevigen/Gibco), we found that both be used at concentrations as low as 2 µg cm^−2^ (a 1:1000 dilution) (Figure S1A) and were subsequently used at a conservative 1:800 dilution for all future experiments.

**Figure 1.**
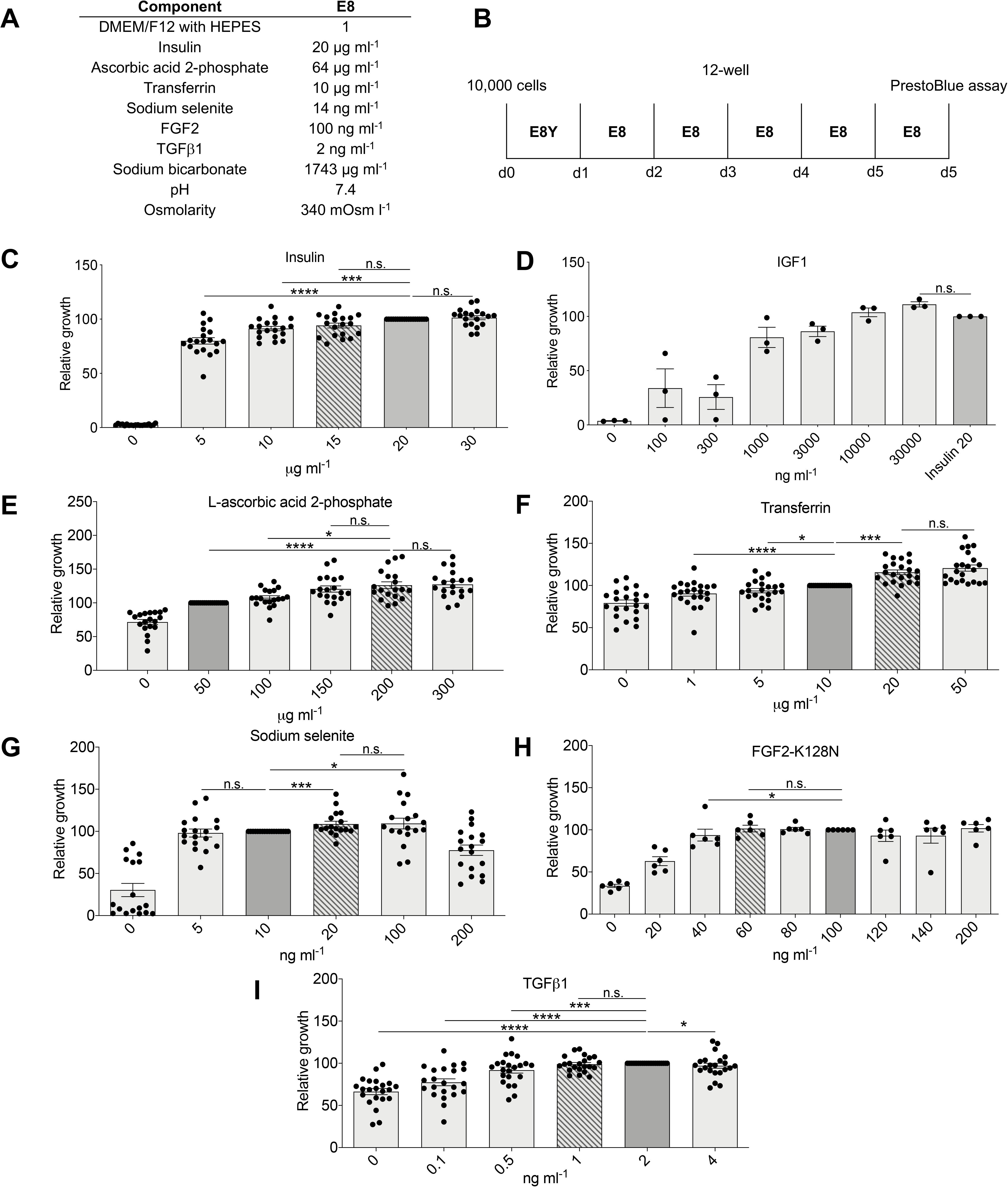
Optimization of Basic Human Pluripotent Stem Cell Medium Constituents with a Short-Term Growth Assay. Results are normalized to initial medium component concentrations shown with a dark gray bar. Optimized component concentrations shown with a diagonal hash. Optimizations were completed using hiPSC line 19c3 and a short-term 6-day growth assay. **(A)** Concentrations of initial media formula components. **(B)** Schematic of assay growth schedule showing seeding of cells at 10,000 cells per well of a 12-well plate, day on which media was changed (E8Y/E8), and the use of a PrestoBlue cell viability assay on day 6 to assess cell number. **(C)** Comparison of the effect on relative growth of concentrations of recombinant human insulin (*n* = 19) **(D)** Recombinant human IGF1 LR3 (*n* = 3). **(E)** L-ascorbic acid 2-phosphate (*n* = 19). **(F)** Transferrin (*n* = 22). **(G)** Sodium selenite (*n* = 18). **(H)** FGF2-K128N (*n* = 6). **(I)** TGFβ1 (*n* = 22). *n* = full experimental replicates, Mann-Whitney test, \**P* ≤ 0.05, \*\**P* ≤ 0.01, \*\*\**P* ≤ 0.005, \*\*\*\**P* ≤ 0.0001, n.s. = not significant. The significance bar refers to the significance between the conditions at the beginning and the end of the bar.

We next established a suitable pluripotent growth assay modeled on that used by Ludwig *et al*. and Chen *et al*. after establishing that neither automated cell counting of dissociated cells (6-well) or small-format plate reader-based cell viability assays (i.e. 96-well) were suitably robust (Figure 1B). This protocol involved dissociating ~75% confluent hiPSC to single cells with TrypLE and seeding 10,000 cells per well of a 12-well plate in medium to be tested in the presence of a Rho kinase inhibitor for 24 h. Medium this then changed every day for 5 days and followed by a PrestoBlue viability assay using a plate reader to assess the number of viable cells present. We utilized this 6-day protocol rather than our typical 4-day protocol to enhance the detection of variance between formulae during sub-optimal low-density culture. All initial experiments were performed using a version of FGF2 we generated in-house with a mutation (K128N) previously demonstrated to enhance thermostability (Chen et al., 2012). In a first round of optimizations we assessed a range of concentrations for each of the six major hiPSC medium components (Figures 1C-I). During this stage we established that insulin was essential, and the effect of insulin was dose-dependent up to 20 µg ml^−1^ (Figure 1C). Insulin could only be replaced by very high levels of IGF1 LR3 (≥1 µg ml^−1^), although this was not cost-effective (Figure 1D). Ascorbic acid 2-phosphate was not essential, as previously demonstrated (Prowse et al., 2010), but higher levels (≥150 µg ml^−1^) enhanced growth (Figure 1E). Of note, this level of ascorbic acid 2-phosphate is similar to the level optimized in the cardiac differentiation media CDM3 (Burridge et al., 2014). Transferrin was also not essential to the media formula, but improved growth in a dose-dependent manner with 20 µg ml^−1^ exhibiting optimal growth while maintaining cost-efficiency (Figure 1G). A source of selenium was shown to be essential, although concentrations of sodium selenite between 5-100 ng ml^−1^ did not significantly affect growth, and sodium selenite became toxic at ≥200 ng ml^−1^ (Figure 1F). FGF2-K128N was optimal at ≥60 ng ml^−1^ (Figure 1H), and TGFβ1 was sufficient at ≥1 ng ml^−1^ in this simple one passage growth assay (Figure 1I).

### Optimization of Additional Media Components

In previous work, Chen *et al*. show that transferrin improves single cell clonality in the absence of the Rho-associated kinase (ROCK1/2) inhibitor Y27632. We could not replicate this observation in our assay (Figure 2A) suggesting that the inclusion of a ROCK1/2 inhibitor for at least the first 24 h after passage is optimal. We also confirmed that the addition of 2 µM thiazovivin for the first 24 h marginally improved growth over 10 µM Y27632 and was ~5 × more cost-effective choice (Figure 2B, 2C). Some recent hiPSC growth formulae have suggested that the addition of high levels (2 ×) of non-essential amino acids (NEAA) and/or low levels (0.1 ×) of chemically defined lipids enhance growth (Figure S1B). In our assay NEAA did not augment growth and the addition of lipids was inhibitory at all but the lowest levels (Figure 2D). Supplementation with bovine serum albumin (BSA), a common hPSC media component, did not have positive or negative effects on growth and was excluded to maintain a chemically defined formula (Figure 2E). The DMEM/F12 basal media we use from Corning contains higher levels of sodium bicarbonate (~29 mM or 2438 mg l^−1^) compared to DMEM/F12 from other manufacturers (Figure S1C). Supplementation of Gibco DMEM/F12, which contains 14 mM of sodium bicarbonate, with 20 mM of additional sodium bicarbonate has recently been demonstrated to be advantageous to hiPSC growth rate by suppressing acidosis of the medium (Liu et al., 2018). In our hands, the standard 29 mM of sodium bicarbonate was optimal (Figure 2F). We also assayed the effect of pH and osmolarity on growth. We found that pH 7.1, measured room temperature and atmospheric CO_2_ and modified with HCl or NaOH (Figure 2G) and a lower osmolarity of 310 mOsm l^−1^, modified NaCl or water (Figure 2H), promoted the highest growth rate in contrast to pH 7.2 and 350 mOsml^−1^ demonstrated by Ludwig *et al*. (Ludwig et al., 2006b) or pH 7.4 and 340 mOsm l^−1^ used by Chen *et al*. (Chen et al., 2011). Of interest here, low osmolarity KnockOut DMEM (270 mOsm l^−1^) has historically been used for mESC culture based on the approximation of mouse embryonic tissue (Lawitts and Biggers, 1992). All of our experiments were optimized for growth at 5% O_2_ and 5% CO_2_, a common compromise for optimal hypoxia which enhances reprograming efficiency (Chen et al., 2011), pluripotency (Forristal et al., 2010), and genome stability (Narva et al., 2013), without the significant cost increase from high N_2_ usage to obtain lower O_2_ levels, or from 10% CO_2_ suggested by some (Ludwig et al., 2006b), that necessitates higher levels of sodium bicarbonate.

**Figure 2.**
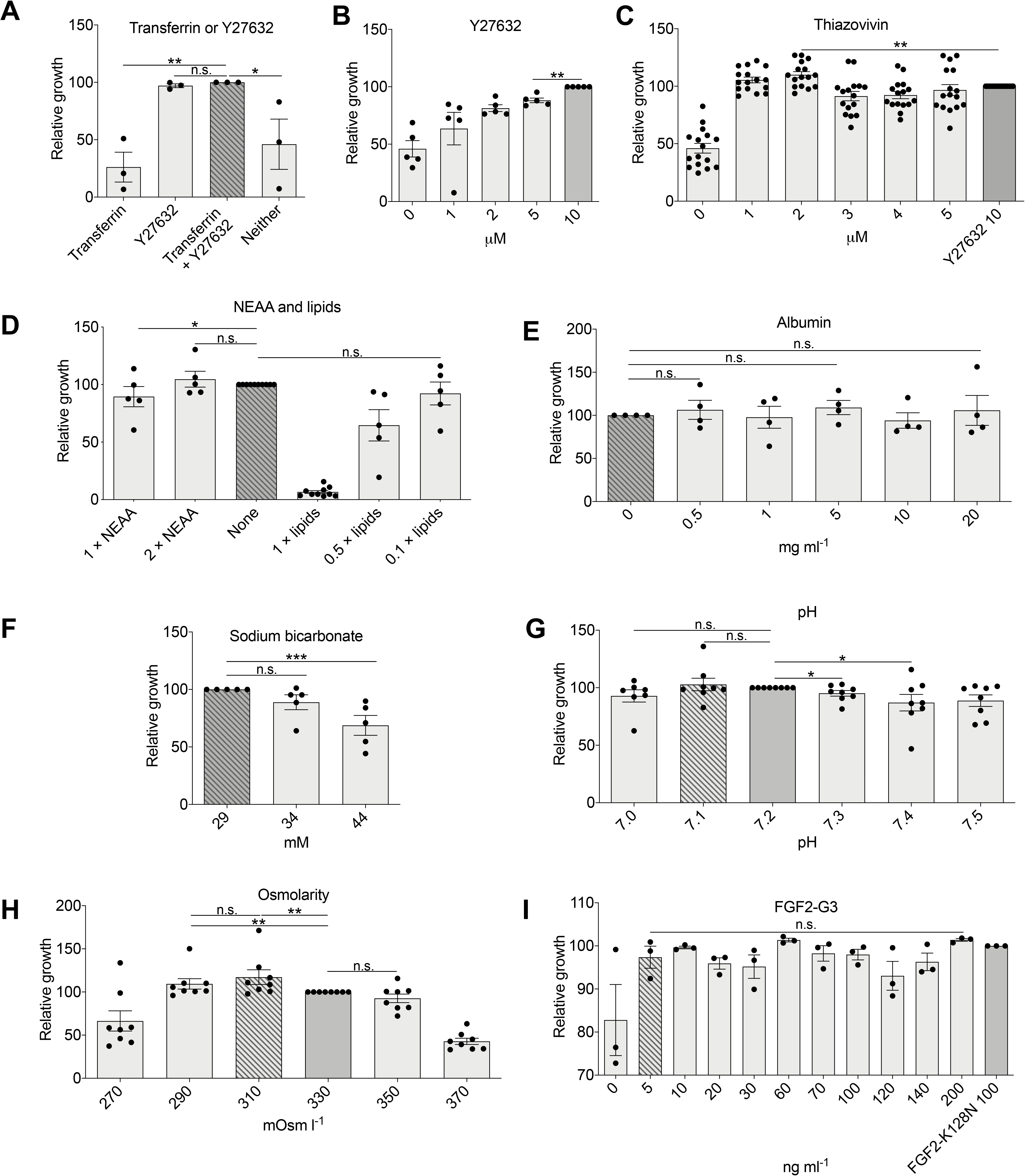
Optimization of Additional Human Pluripotent Stem Cell Medium Constituents in a Short-Term Assay. Results are normalized to initial medium component concentrations shown with a dark gray bar. Optimized component concentrations shown with a diagonal hash. Optimizations were completed using hiPSC line 19c3 and a short-term 6-day growth assay. **(A)** Comparison of the suitability of recombinant transferrin (10 µg ml^−1^) to support clonal growth with and without ROCK1/2 inhibition using Y27632 (10 µM) during the first 24 h after passage (*n* = 3). **(B)** Comparison of ROCK1/2 inhibitor Y27632 only during first 24 h after passage on relative growth (*n* = 5). **(C)** Comparison of ROCK1/2 inhibitor thiazovivin only during first 24 h after passage on relative growth (*n* = 16). **(D)** Comparison of the effect of the addition of non-essential amino acids (NEAA) and chemically defined lipids (*n* = *5*) on relative growth. **(D)** Fatty acid-free albumin (*n =* 4). **(E)** Sodium bicarbonate (*n* = 5). **(F)** pH (*n* = 9). **(G)** Osmolarity (*n* = 8). **(H)** FGF2-G3 (*n* = 3). *n* = full experimental replicates, Mann-Whitney test, \**P* ≤ 0.05, \*\**P* ≤ 0.01, \*\*\**P* ≤ 0.005, \*\*\*\**P* ≤ 0.0001, n.s. = not significant. The significance bar refers to the significance between the conditions at the beginning and the end of the bar.

In our original hiPSC culture medium formula, we used commercial recombinant human FGF2, and this accounted for >60% of the media cost. This led us to investigate the production of FGF2 in-house, for which we generated a FGF2 sequence with *E. coli*-optimized codon usage to enhance yield and a K128N point mutation to improve thermostability (Chen et al., 2012). This sequence was inserted into a recombinant protein production plasmid (pET-28a) which utilizes an upstream 6×His tag for purification that was not cleaved during processing (Figure S2). We subsequently generated two additional thermostable FGF2 variants: FGF1-4X (Chen et al., 2012), and FGF2-G3 (Dvorak et al., 2018). We found FGF2-G3, with nine point-mutations was more potent than FGF2-K128N, showing a similar effect on growth rate at 5 ng ml^−1^ to FGF2-K128N at 100 ng ml^−1^ (Figure 2I) when tested using this short-term assay.

### Optimization of Media Components using a Long-Term Assay

Our initial short-term experiments were useful for preliminary optimizations, yet we were aware from previous experiments that some variables, such as the elimination of TGFβ1, had minimal effects short-term and would only have detectable negative effects in long-term experiments. We began a second round of optimizations using a long-term version of our growth assay covering 5 passages (Figure 3A). Each experiment was independently repeated at least 5 times. These experiments again confirmed optimal concentrations of insulin (20 µg ml^−1^; Figure 3B), ascorbic acid 2-phosphate (200 µg ml^−^1; Figure 3C), transferrin (20 µg ml^−1^; Figure 3D), sodium selenite (20 ng ml^−^1; Figure 3E), and FGF2-G3 (40 ng ml^−1^; Figure 3F).

**Figure 3.**
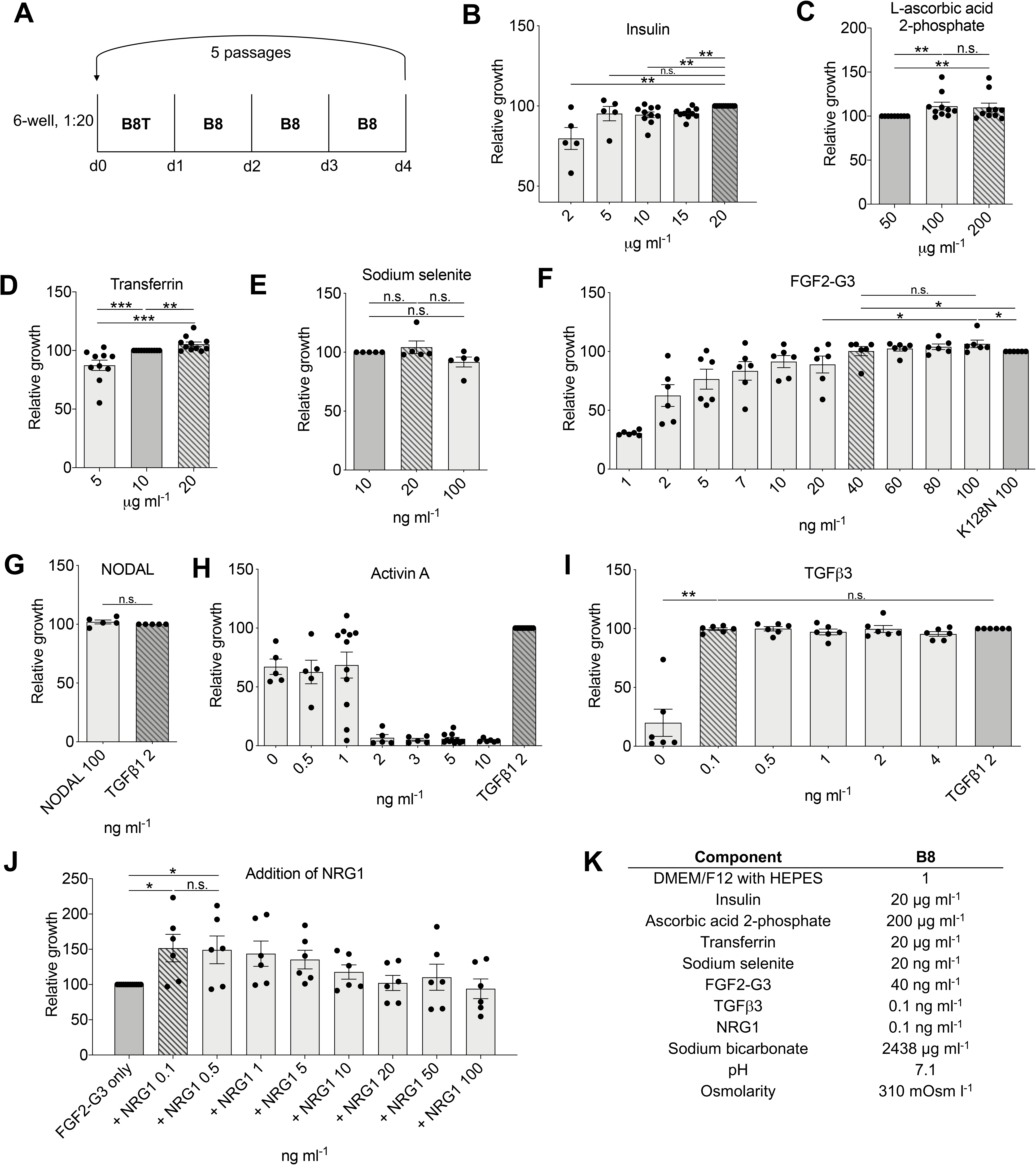
Optimization of B8 Medium Constituents with a Long-Term Growth Assay. Results are normalized to initial medium component concentrations shown with a dark gray bar. Optimized component concentrations shown with a diagonal hash. Optimizations were completed using in hiPSC line 19c3 using a long-term 5-passage, 4-day growth assay. Optimized component concentrations shown with a diagonal hash. **(A)** Schematic of passaging schedule. **(B)** Comparison of the effect of recombinant human insulin concentration on relative growth (*n* = 10). **(C)** L-ascorbic acid 2-phosphate (*n* = 13). **(D)** Recombinant transferrin (*n =* 10). **(E)** Sodium selenite (*n = 5*). **(F)** In-house made FGF2-G3 (*n* = 6). **(G)** NODAL (*n* = 5). **(H)** Activin A (*n* = 5). **(I)** In-house made TGFβ3 after 9 passages compared to commercial TGFβ1 (*n* = 9). **(J)** Addition of NRG1 to 40 ng ml^−1^ FGF2-G3 (*n* = 6). **(K)** Final B8 formula. *n* = full experimental replicates, Mann-Whitney test, \**P* ≤ 0.05, \*\**P* ≤ 0.01, \*\*\**P* ≤ 0.005, \*\*\*\**P* ≤ 0.0001, n.s. = not significant. The significance bar refers to the significance between the conditions at the beginning and the end of the bar.

We next looked for alternatives to TGFβ1 as this was now the costliest component (~40% of total cost) and found that this could be not used at lower concentrations than previously suggested (2 ng ml^−1^) when using our long-term assay (Figure S3A). We found that other Activin/NODAL/TGFβ1 signaling sources such as NODAL were required at cost-prohibitively high levels (100 ng ml^−1^) (Figure 3G), and Activin A was not suitable to maintain growth to the same level as TGFβ1 (Figure 3H). Activin A in combination with TGFβ1 also has had a negative effect on growth (Figure S3B). TGFβ1 is a homodimer of two *TGFB1* gene products and therefore the recombinant protein is commonly produced in mammalian cells making it complex to produce for basic research labs. These mammalian cells provide post-translational modifications crucial for biological activity such as glycosylation, phosphorylation, proteolytic processing, and formation of disulfide bonds which are not present in *E. coli*. To overcome this issue, we generated a version of TGFβ1 (TGFβ1m) that is unable to form dimers due to a point mutation. This monomeric protein is predicted to be ~20-fold less potent than TGFβ1 but can be easily produced in large quantities in *E. coli* (Kim et al., 2015). Our initial experiments demonstrated the *TGFB1* sequence with *E. coli*-optimized codon usage was expressed in inclusion bodies. To overcome this, we inserted the sequence into a protein production plasmid (pET-32a) designed to produce a thioredoxin fusion protein allowing expression in the cytoplasm. It has been also demonstrated that TGFβ3 is more potent than TGFβ1 (Huang et al., 2014). Comparing TGFβ1, TGFβ1m, TGFβ3, and TGFβ3m, we found that TGFβ3 offered the best combination of being able to be produced in *E. coli* and suitable for use at 0.1 ng ml^−1^ (Figures 3I, S3C, and S3D) and was therefore selected for the final formula.

Finally, two hESC media formulations that we studied, DC-HAIF and iDEAL, contain both FGF2 and neuregulin 1 (NRG1) (Figure S1B). We found that supplementation with all tested levels of NRG1 enhanced growth by >15% over FGF2-G3 alone (Figures 3J and S3E), although NRG1 is not able to support growth in the absence of FGF2 (Figure S3F). We similarly generated recombinant NRG1 protein using pET-32a, which prevented production as inclusion bodies, with subsequent cleavage using thrombin to form an active protein (Figures S3G and S4). Our final B8 media formulation was derived from these long-term assay optimizations and is shown in Figure 3K.

### Demonstration of the Suitability of B8 for hiPSC Generation and Weekend-Free Culture

The gold-standard demonstration of the suitability of a hiPSC media is the capacity to generate hiPSC lines and maintain them in long-term culture. Our standard hiPSC line 19c3 has been cultured for >100 passages in B8 (Figure 4A) and maintained expression of markers of undifferentiated status (SSEA4 and TRA-1-60) by flow cytometry (p131 at time of assay). We also generated hiPSC lines from 37 patients using established protocols but using B8. These hiPSC lines maintained SSEA4 and TRA-1-60 expression in culture long-term (up to p65 at time of assay) (Figure 4A). In B8 hiPSC maintained a hESC-like morphology (Figure 4B), positive immunofluorescent staining for SSEA4, POU5F1, SOX2, and TRA-1-60 (Figure 4C), and normal karyotype (Figure 4D). We then compared this B8 formula (Figure 3K) with our traditional E8 formula and found that growth similar to B8 across commonly used split ratios (Figure 4E). It has been demonstrated that unlike commercial FGF2, thermostable variants of FGF2 such as FGF2-G3 are capable of inducing pERK in FGF-starved cells, even after media had previously been stored for extended periods at 37 °C (Chen et al., 2012; Dvorak et al., 2018). To confirm that our FGF2-G3 performed similarly we performed a comparable assay and corroborated that FGF2-G3 was stable after 7 days at 37 °C, whereas commercial FGF2 was not capable of stimulating pERK after 2 days at 37 °C. As common ‘weekend-free’ media formulae such as StemFlex have reverted to using albumin and likely contain heparin, both of which are not chemically defined, we also assessed the function of these components in this assay and found that both improved the stability of FGF2-G3 (Figure 4F).

**Figure 4.**
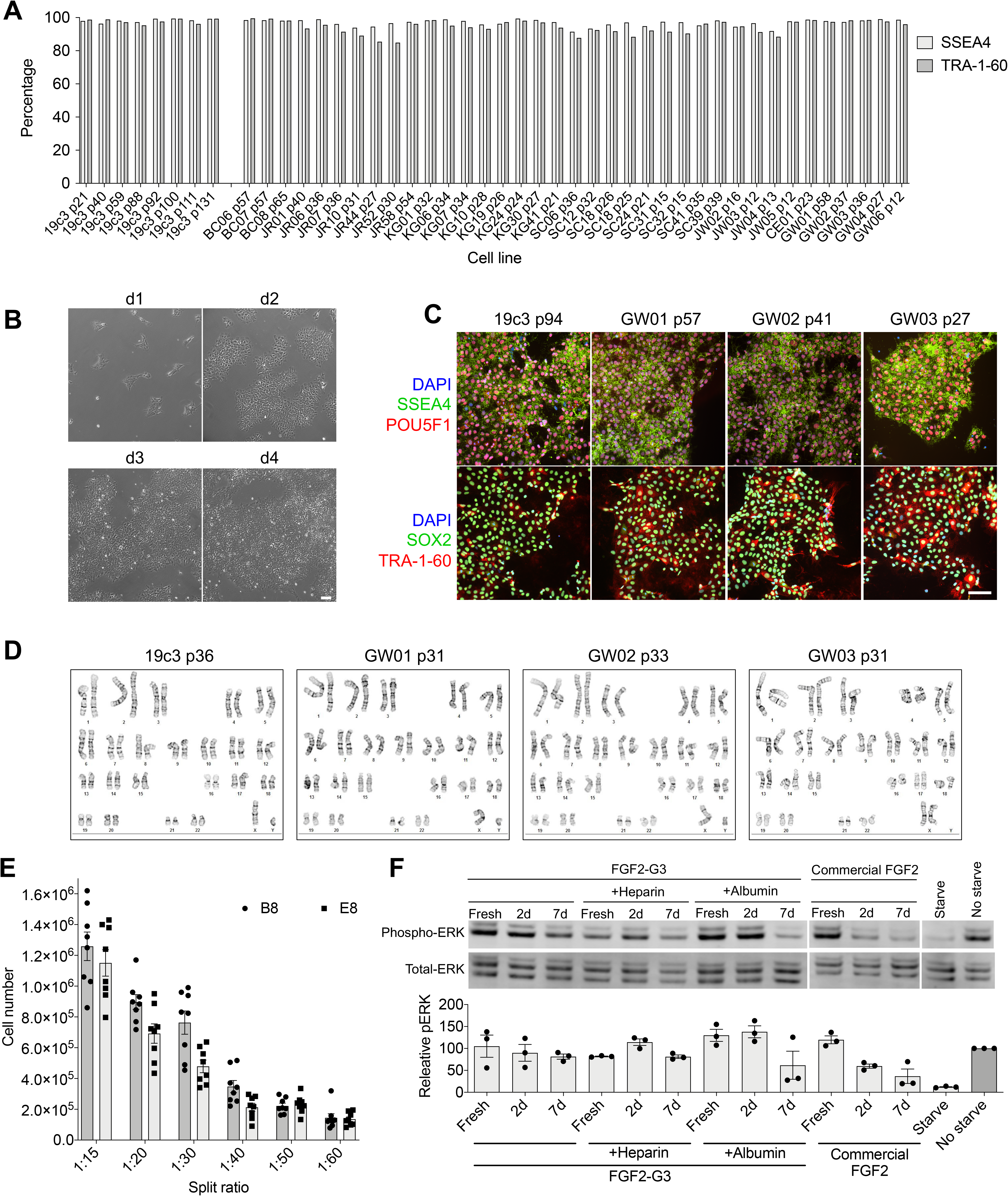
Qualification of B8 as Suitable for hiPSC Generation and Culture. **(A)** Demonstration of expression of markers of undifferentiated status assessed by flow cytometry in hiPSC line 19c3 cultured in B8 from passage (p) 21 to 131 (left) and 37 individual hiPSC lines derived in B8 from passage 12 to 65 (right) all derived in B8. **(B)** Phase contrast images (10x) of hiPSC line 19c3 p44 cultured in B8. Scale bar, 100 µm **(C)** Expression of markers of pluripotency and undifferentiated status in a variety of B8-derived hiPSC lines. Scale bar, 100 µm **(D)** Example G-banding karyotype analysis of four hiPSC lines derived in B8. **(E)** 19-3 hiPSC growth at low seeding densities in B8 compared to E8 (*n* = 8). **(F)** Assessment of stimulation of phospho-ERK after media had been stored at 37 ºC for 2 or 7 days, comparing in-house generated FGF2-G3 to a commercial FGF2 (Peprotech). hiPSC were starved of FGF2 for 24 h then treated with the indicated media for 1 h before collection for Western blot. Total ERK was used as a loading control.

With the understanding that B8 supports hiPSC growth across a variety of sub-optimal conditions (Figure 4E), we established the suitability of this formula to skip days of media change. We established a matrix of media change days both with and without thiazovivin (Figure 5A) and showed that daily B8 media changes (top row) was surprisingly one the least suitable for growth rate. B8T treatment without media changes was the least effective of the thiazovivin-containing protocols (bottom row), whereas B8T followed by B8 (dark grey bar) was a suitable compromise between growth rate and extended exposure to thiazovivin. With the knowledge of the influence of various media change-skipping timelines, we devised two 7-day schedules consisting of passaging cells every 3.5 days that would allow for the skipping of media changes on the weekends, a major caveat of current hiPSC culture protocols (Figures 5B and 5C). Our data demonstrated that a weekend-free 3.5-day schedule with medium exchange after the first 24 h was suitable long-term, whereas no media exchange (i.e. passage only) was less suitable over a 25-passage timeline (Figure 5D). In addition, supplementation of the media with 0.5 mg ml^−1^ of albumin did not rescue this deficit (Figure 5D). Further experiments with various doses of albumin or heparin further confirmed that these do not have an additive effect on weekend-free growth in B8 (Figure 5E). One of the major caveats of skipping media changes is that differentiation efficiency and reproducibility is diminished. We used our existing cardiac (Burridge et al., 2014), endothelial (Patsch et al., 2015), and epithelial differentiation (Li et al., 2015; Qu et al., 2016) protocols to demonstrated that B8 supported differentiation to a similarly high level with either daily media change or our weekend-free protocol (Figures 5F, 5G, 5H).

**Figure 5.**
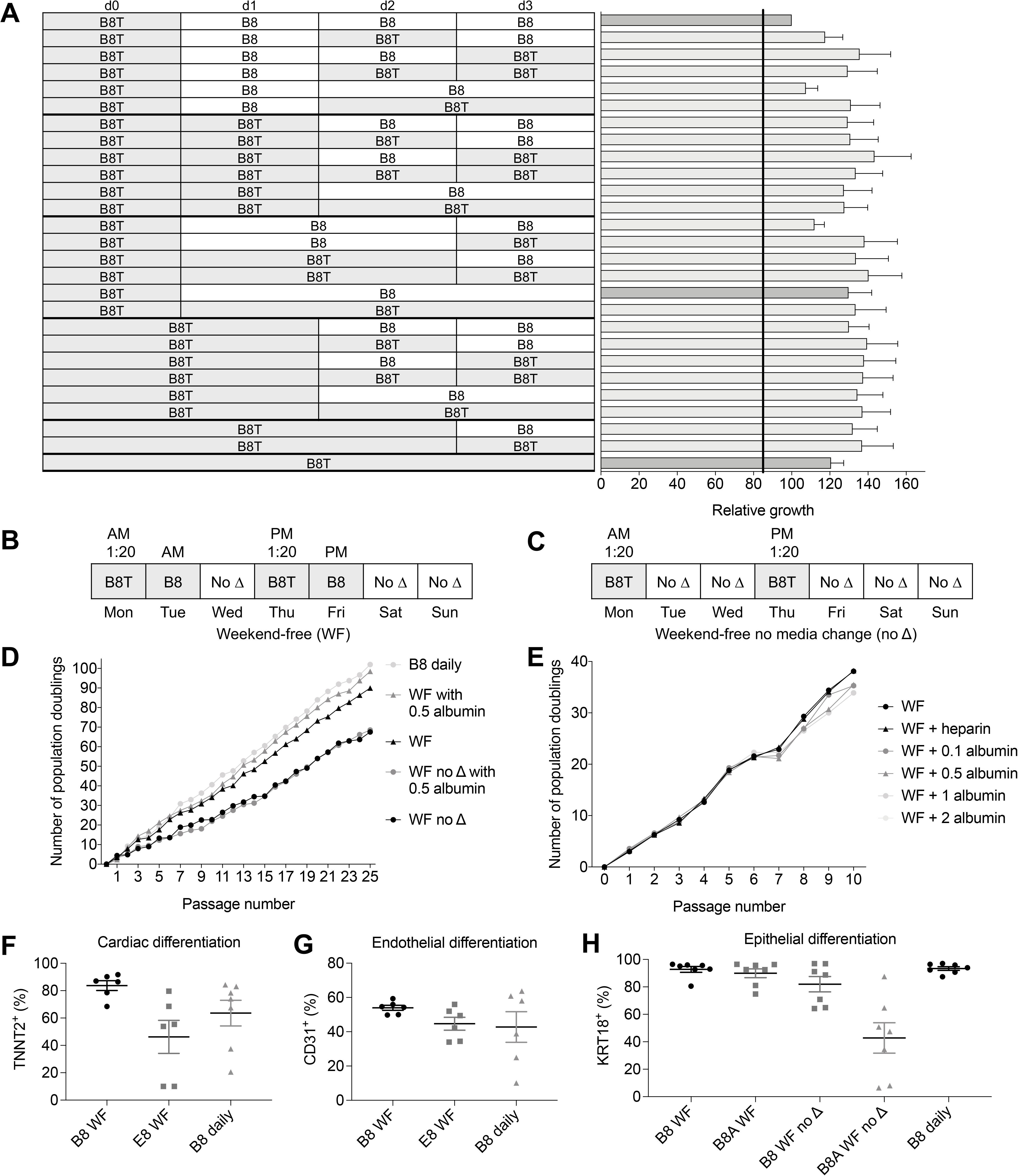
Optimization of Weekend-Free Passaging Schedule that is Still Compatible with Monolayer Differentiation. **(A)** Establishment of an optimal 4-day media change schedule. Experiments were completed with hiPSC line 19c3. Grey box represents B8T treatment, white box represents B8 treatment, width of box represents number of days of culture in that media (*n* = 8). **(B)** Weekend-free (WF) passage and media change schedule. Grey boxes represent days of media change. **(C)** Weekend-free schedule without media change (WF no Δ) schedule. **(D)** Comparison of growth when using the WF and WF no Δ schedules, with or without addition of 0.5 mg ml^−1^ albumin over 25 passages. **(E)** Comparison the addition of varying levels of albumin (mg ml^−1^) to a WF schedule (*n* = 4). **(F)** Cardiac differentiation efficiency when using WF schedule (*n* = 5). **(G)** Endothelial differentiation efficiency when using WF schedule (*n* = 6). (H) **(G)** Epithelial differentiation efficiency when using WF schedule with or without addition of 0.5 mg ml^−1^ albumin (A) (*n* = 7).

## Discussion

The hiPSC field predominantly utilizes two hiPSC culture media, the commercial mTeSR1 and Essential-8. Despite the formulae for these media being available to researchers, the complexity of media compounding and cost of growth factors required have made this unappealing. These two media, along with six other published hiPSC culture media formulae, have much in common due to their shared history (Figure S1B). Despite this, there are many media components that have not had their concentration optimized or necessity demonstrated. In many cases, especially the formulation of TeSR, optimization experiments were discussed yet no data were provided. In the experiments here we found that only five components were truly essential for hiPSC culture: insulin, sodium selenite, FGF2, DMEM/F12, (Figure 1) and importance of a fifth component TGFβ1 was only evident in the long-term assay (Figures 1I, 3I, and S3A, C, D). The other three components, ascorbic acid 2-phosphate, transferrin, and NRG1, are dispensable for hiPSC growth, although their removal results in a reduced growth rate.

During the development of this assay platform, we found that precise measurement of relative growth was a suitable surrogate for metrics of the pluripotent state (such as *NANOG* expression). When cells began to spontaneously differentiate, such as when TGFβ1 is omitted from the formula, the resulting slowing of growth was easily detectable. We noted a number of surprising results during this optimization. For example, we could not find a positive or negative effect of the addition of albumin (Figures 2D, 5D) despite it being a common constituent of many academic and commercial media formulae. We propose that high levels of ascorbic acid in our formula may have eliminated some of the need for the antioxidant role of albumin. We found that Activin A was not suitable either with or without TGFβ1 (Figure 3H and S3B) despite its inclusion in a variety of other formulae. We do show that at very low doses Activin A could support growth, albeit to a lesser extent than TGFβ1. In some media formulae, such as hESF9T, there is a positive correlation between higher levels of TGFβ1 (up to 10 ng ml^−1^) and expression of *NANOG* (Yamasaki et al., 2014) while others have found only minimal differences above 0.1 ng ml^−1^ (Frank et al., 2012). Similarly, it was shown that higher levels of HSA correlated with higher levels of *POU5F1*, *SOX2*, and *NANOG* (Frank et al., 2012) yet albumin-free formulae have been used extensively without issue. One of our major efforts during the development of B8 was the minimization of media costs.

Typical list prices of 1 liter of commercial media such as Essential 8 or mTeSR1 are $450-550 USD (Figure S5A) plus shipping. In our previous E8-based formula, the growth factors FGF2 and TGFβ1 represented more than 80% of the total medium costs and we found that this could be much higher if laboratories could not negotiate discounts from commercial recombinant protein suppliers resulting in a cost of ~$115 per liter (Figure S5A). The generation of recombinant proteins in-house, shown here to be major factor in reducing media costs, although daunting, was highly simplified by using three commercial products: MagicMedia for *E. coli* growth, B-PER for lysis, and Ni-NTA spin columns for purification. Combined, these eliminate much of the complexity for recombinant protein production, including the need for HPLC or FPLC. Each of these components could easily be replaced by more cost-effective procedures such as inducing protein expression using IPTG, making lysis buffers, or using higher-throughput columns, albeit with greater complexity. Our optimization of the plasmids and generation of thioredoxin fusion proteins where necessary eliminates much of the complexity associated with inclusion bodies and the resulting refolding processes otherwise required. A typical 1 liter *E. coli* culture, which requires two days and basic laboratory skills, will usually provide 80 mg of FGF2-G3, enough for 800 liters of B8. Similarly, a 500 ml culture of TGFβ3 or NRG1 will commonly provide enough protein years of work (~800,000 liters of B8 media). This optimization results in a media cost of ~$16 per liter and leaves insulin and transferrin as the major remaining costs, although the concentrations of these components could also potentially be reduced with only minor impact on growth rate (Figure 3B, 3D) reducing costs to ~$12 per liter. The difference between growth rate of hiPSC in either formula is likely undetectable in normal culture. All costs described do not include shipping or technician time for compounding and quality control, which limit savings for labs that use very small quantities of B8. This issue may be overcome by multiple laboratories pooling resources or an institute’s core facility taking responsibility for supplement aliquot generation.

The final major cost is the basal media DMEM/F12, now representing ~75% of the cost of the B8 formula (Figure S5A). It is simple to generate DMEM/F12 from powder to reduce costs, although the requirement for suitable water quality and maintaining these systems, and the use of filter sterilization adds cost and complexity of the media making process and therefore may not be appropriate for all but the highest usage labs. DMEM/F12 is a combination of 52 amino acids, vitamins, inorganic salts, and other components derived from DMEM, a high nutrient (amino acid and vitamin) concentration media, originally optimized for fibroblast growth, and F12, a rich and complex fatty acid-containing media optimized for CHO cells to generate a complex ‘catch-all’ formula. Few experiments have been completed to compare alternatives to DMEM/F12 for hiPSC culture, indeed, Chen *et al*. showed comparable results between DMEM/F12 and the comparatively simple MEMα, suggesting that there is much to learn in this respect.

hiPSC culture without daily media changes introduces a number of potential caveats, for example, we know that the L-glutamine in the medium is unstable at 37 °C and that the concentration is reduced by about a third over four days. We also know that cells are producing lactate and ammonia, reducing the pH of the media, although this buffered partially by the HEPES and sodium bicarbonate. Finally, hiPSCs release autocrine or paracrine factors into the media that may induce differentiation and the increase in these factors over time has not been decoupled from the use of nutrients and production of metabolic waste. We chose not to maintain exposure to Rho-associated kinase (ROCK1/2) inhibitors as there exists data suggesting that this induces epithelial-to-mesenchymal transition (Maldonado et al., 2016). Furthermore, ROCK inhibitors have a wide-range of off-target effects (Andrews et al., 2010). These issues may result in undesired effects although this remains to be validated experimentally and formulations do exist that maintain constant ROCK1/2 inhibitor exposure (Tsutsui et al., 2011).

As hiPSC projects become larger and move beyond proof-of-principle experiments, the labor required to maintain cells becomes burdensome, especially the need for daily media changes 7-days a week. Here we show that our B8 formula is well-suited to weekend-free schedules. A major issue with some commercial media is that although a weekend-free schedule is feasible, growth of hiPSCs is considerably slower and it is recommended to grow cells as low-density colonies. These low-density colonies are not compatible with subsequent monolayer differentiation protocols, as have become commonplace with the majority of lineages. The optimization of B8 specifically for fast monolayer growth, along with the incorporation of thermostable FGF2-G3, overcomes many of these issues while maintaining compatibility with common differentiation protocols and has become standard practice within our laboratory.

## Experimental Procedures

### Human Induced Pluripotent Cell Culture

All pluripotent and reprogramming cell cultures were maintained at 37 ºC in Heracell VIOS 160i copper-lined direct heat humidified incubators (Thermo Scientific, 51030410) with 5% CO_2_ and 5% O_2_. N_2_ for incubators was derived from the gas port of a 100 psi, 230-liter liquid nitrogen tank. Differentiation cultures were maintained at 5% CO_2_ and atmospheric (~21%) O_2_. All cultures (pluripotent and differentiation) were maintained with 2 ml medium per 9.6 cm^2^ of surface area or equivalent. All media was used at 4 °C and was not warmed to 37 °C before adding to cells due to concerns of the thermostability of the FGF2 (Chen et al., 2012). We have found no detectable effects on cell growth from using cold media. All cultures were routinely tested for mycoplasma using a MycoAlert PLUS Kit (Lonza) and a 384-well Varioskan LUX (Thermo Scientific) plate reader. E8 medium was made in-house as previously described (Burridge et al., 2015; Chen et al., 2011) and consisted of DMEM/F12 (Corning, 10-092-CM), 20 μg ml^−1^ *E. coli*-derived recombinant human insulin (Gibco, A11382IJ), 64 μg ml^−1^ L -ascorbic acid 2-phosphate trisodium salt (Wako, 321-44823), 10 μg ml^−1^ *Oryza sativa*-derived recombinant human transferrin (Optiferrin, InVitria, 777TRF029-10G), 14 ng ml^−1^ sodium selenite (Sigma, S5261), 100 ng ml^−1^ recombinant human FGF2-K128N (made in-house, see below), 2 ng ml^−1^ recombinant human TGFβ1 (112 amino acid, HEK293-derived, Peprotech, 100-21). Cells were routinely maintained in E8 medium on 1:800 diluted growth factor reduced Matrigel (see below). E8 was supplemented with 10 μM Y27632 dihydrochloride (LC Labs, Y-5301), hereafter referred to as E8Y, for the first 24 h after passage. For standard culture, cells were passaged at a ratio of 1:20 every 4 days after achieving ~70-80% confluence using 0.5 mM EDTA (Invitrogen, 15575020) in DPBS (without Ca^2+^ and Mg^2+^, Corning, 21-031-CV). Cell lines were used between passages 20 and 100. Other media components tested were: Human Long R3 IGF1 (Sigma, 91590C), thiazovivin (LC Labs, T-9753), recombinant human TGFβ3 (Cell Guidance Systems, GFH109), sodium bicarbonate (Sigma, S5761), NEAA (Gibco, 11140050), Chemically Defined Lipid Concentrate (Gibco, 11905031), fatty acid-free bovine serum albumin (GenDEPOT, A0100). pH was adjusted with 1N HCl (Sigma, H9892) or 1N NaOH (Sigma, S2770) and measured at room temperature and atmospheric CO_2_ using a SevenCompact pH meter (MettlerToledo). Osmolarity was adjusted with sodium chloride (Sigma, S5886) or cell culture water (Corning, 25-055-CV) and measured with a Vapro 5600 vapor pressure osmometer (Wescor).

### Matrigel Optimization

Our standard condition throughout was 2 ml of 1:800 reduced growth factor Matrigel (Corning, 354230) diluted in 2 ml of DMEM (Corning, 10-017-CV) per well of 6-well plate (Greiner, 657165) or equivalent. Geltrex (Gibco, A1413201), Cultrex (Trevigen, 3433-005-01), and Synthemax II-SC (3535, using a 1:320 dilution) were tested as previously detailed (Burridge et al., 2014) and found to be successful for growth. Plates were made and kept in CO_2_ incubators at 37 °C for up to one month. A variety of Matrigel lot numbers were used throughout this project with concentrations between 9.7 and 10.8 mg ml^−1^. For the concentration optimization experiments (Figure S1A) lot number 8162008 (10.7 mg ml^−1^) was used.

### FGF2-K128N Generation

The full length (154 amino acid) FGF2 sequence: AAGSITTLP ALPEDGGSGA FPPGHFKDPK RLYCKNGGFF LRIHPDGRVD GVREKSDPHI KLQLQAEERG VVSIKGVCAN RYLAMKEDGR LLASKCVTDE CFFFERLESN NYNTYRSRKY TSWYVAL<u>**K**</u>RT GQYKLGSKTG PGQKAILFLP MSAKS with a K128N substitution (in bold/underlined) was codon optimized for *E. coli* with the addition of a *Bam*HI site at the start (5’) of the sequence and an *Eco*RI site at the end (3’). This sequence was synthesized on a BioXp 3200 (Synthetic Genomics). The insert was then digested with *Bam*HI and *Eco*RI (Anza, Invitrogen) and ligated with T4 DNA ligase (Anza) into a pET-28a expression vector (Novagen, 69864) and cloned in to One Shot BL21 Star (DE3) chemically competent *E. coli* (Invitrogen, C602003)*. E. coli* were stored in 25% glycerol (Invitrogen, 15514011) at −80 °C. A starter cultured was prepared by inoculating 5 ml of LB (Fisher, BP1425-500) supplemented with 50 µg ml^−1^ kanamycin sulfate (Gibco, 11815024) in a bacterial tube (Corning Falcon) and incubated in an Innova-44 Incubator-Shaker (New Brunswick) at 220 rpm overnight at 37 °C. Protein expression was performed in a 2000 ml baffled shaker flask (Fisherbrand, BBV2000) as follows: The 5 ml of starter culture was added to 500 ml of MagicMedia (Invitrogen, K6815), supplemented with 50 µg/ml kanamycin sulfate and incubated as above for 24 h at 30 °C. The culture was harvested in to 2 × 250 ml centrifuge bottles (Nalgene, 3120-0250) and centrifuged in a J6-MI centrifuge (Beckman Coulter) at 3,000 rpm for 30 min at 4 °C. Supernatant was carefully poured off and pellets were weighed and stored at −80 °C for downstream processing. Cell pellets were resuspended B-PER lysis buffer (Thermo Scientific, 78248) using 5 ml of B-PER Complete Reagent per gram of bacterial cell pellet, neither DNase nor lysozyme were added. Cells were incubated for 15 minutes at RT with gentle rocking. The bottles containing the lysates were then centrifuged in an ultracentrifuge at 16,000 × *g* for 20 min at 4 °C. Supernatants were collected and the cell debris was discarded. Purification was completed using a 3 ml HisPur Ni-NTA Spin Purification Kit (Thermo Scientific, 88229) following the manufacturers recommendations. To enhance the protein binding efficiency to the resin bed, the sample was incubated for 30 min at 4 °C. Four elutes were collected, one every 10 min. The columns were reused following the manufacturer’s regeneration protocol. The protein concentration was evaluated using Quant-iT Qubit Protein Assay Kit (Invitrogen, Q3321) on a Qubit 3 fluorometer. The 6xHis tag was not cleaved as it has been previously demonstrated to not interfere with the FGF2 function (Soleyman et al., 2016). A standard 1 liter culture produced 80 mg of FGF2. Protein in elution buffer was separated into 4 mg aliquots and stored at −20 °C. No concentration or buffer exchange to remove imidizole was required.

### FGF2-G3 Generation

154 amino acid sequence: AAGSITTLP ALPEDGGSGA FPPGHFKDPK <u>**L**</u>LYCKNGGFF LRIHPDGRVD G<u>**T**</u>R<u>**D**</u>KSDP<u>**F**</u>I KLQLQAEERG VVSIKGVCAN RYLAMKEDGR L<u>**Y**</u>A<u>**I**</u>K<u>**N**</u>VTDE CFFFERLE<u>**E**</u>N NYNTYRSRKY<u>**P**</u>SWYVALKRT GQYKLGSKTG PGQKAILFLP MSAKS with R31L, V52T, E54D, H59F, L92Y, S94I, C96N, S109E, and T121P substitutions (in bold) was codon optimized for *E. coli* was generated as above. FGF1-4X was similarly generated.

### TGFβ3 Generation

112 amino acid sequence: ALDTNYCFRN LEENCCVRPL YIDFRQDLGW KWVHEPKGYY ANFCSGPCPY LRSADTTHST VLGLYNT LNP EASASPCCVP QDLEPLTILY YVGRTPKVEQ LSNMVVKSCK CS was codon optimized for *E. coli* and generated as above, ligated into a pET-32a expression vector, and cloned in to One Shot BL21 Star (DE3) with selection using 100 µg ml^−1^ ampicillin (Gibco, 11593027). Protein expression was completed at 30 °C. The use of pET-32a results in the production of a thioredoxin-TGFBβ3 fusion protein which prevents protein expression in inclusion bodies. It is not necessary to cleave the thioredoxin for TGFBβ3 to be active. TGFβ1, TGFβ1m (C77S), and TGFβ3m (C77S) were similarly generated.

### NRG1 Generation

65 amino acid sequence: SHLVKCAEKE KTFCVNGGEC FMVKDLSNPS RYLCKCPNEF TGDRCQNYVM ASFYKHLGIE FMEAE, which is a truncated version of NRG1 containing just the EGF domain, was codon optimized for *E. coli* and generated as above, ligated into a pET-32a (100 µg ml^−1^ ampicillin) expression vector, and cloned in to One Shot BL21 Star (DE3) with selection using ampicillin. Protein expression was completed at 37 °C. The use of pET-32a results in the production of a thioredoxin-NRG1 fusion protein which prevents protein expression in inclusion bodies. It was found necessary to cleave the thioredoxin using Thrombin CleanCleave Kit (Sigma, RECOMT), followed by repurification, keeping the supernatant.

### Media Variable Optimization Protocol

hiPSC line 19c3 (p20-p60), derived from a healthy male was used for media variable optimization. Prior to assay, cells were grown to 75% confluence after 4 days of culture as above. Cells were dissociated with TrypLE (Gibco, 12604013) for 3 min at 37 **°**C and cells were resuspended in DMEM/F12, transferred to a 15 ml conical tube (Falcon) and centrifuged at 200 × *g* for 3 min (Sorvall ST40). The pellet was resuspended in DMEM/F12, diluted to 1 × 10^5^ cells per ml and 10,000 cells were plated per well in Matrigel (1:800)-coated 12-well plates (Greiner) in the medium to be tested along with 2 µM thiazovivin for the first 24 h. Media were changed daily and cells were grown for 6 days. This lower than normal seeding density was used to allow the discovery of factors only detectable under more extreme conditions and therefore provide data on the robustness to the formulation. Cell growth was then assessed using PrestoBlue (Invitrogen, A13262), using 2 h of incubation, and fluorescence (560 nm excitation, 590 nm emission) was measured using a Varioskan LUX (Thermo Scientific) plate reader.

### Western Blot

Stock lysis buffer was prepared as 150 mM NaCl, 20 mM Tris pH 7.5, 1 mM EDTA, 1 mM EGTA, and 1% Triton X-100 and stored at 4 ºC. Fresh complete lysis buffer was prepared with final concentration of 1× Protease Inhibitor (Roche, 5892791001), 1× Phosphatase Inhibitor Cocktail 2 (P5726, Sigma), 1× Phosphatase Inhibitor Cocktail 3 (Sigma, P0044), 2 mM PMSF (Sigma, P7626), and 1% SDS Solution (Fisher, BP2436200). hiPSC were starved using B8 without FGF2 for 24 hours, then treated with media containing the corresponding FGF2 for 1 h. Media was then removed, and cells were washed once with DPBS, harvested with 0.5 mM EDTA in DPBS, and transferred into tubes. Samples were pelleted by centrifugation at 500 × *g* for 3 minutes and the supernatant was discarded. The pellet was resuspended in 150 μl complete lysis buffer and incubated on ice for 30 min. Clear lysates were collected by centrifugation at 10,000 × *g* for 10 min at 4 ºC. The protein concentration was measured with Qubit Protein Assay Kit (Invitrogen, Q33211) and Qubit 4 fluorometer. Lysates was stored in −80 ºC before use. 10 μg of sample was prepared with NuPAGE LDS Sample Buffer (Invitrogen, B0007) and NuPAGE Reducing Agent (Invitrogen, B0009) according to the manufacturer’s instructions and run on NuPAGE 10% Bis-Tris Gel (Invitrogen, NP0302BOX) and Mini Gel Tank system (Invitrogen, A25977) with Bolt MES SDS Running Buffer (Invitrogen, B000202) at 100 V for 1 h. SeeBlue Plus2 Pre-Stained Protein Standard (Invitrogen, LC5925) was used as a ladder. The gel was then transferred in Mini Trans-Blot Cell system (Bio-Rad, 1703930) on to a PVDF transfer membrane (Thermo Scientific, 88518) at 240 mA for 1 h and 30 min. The membrane was blocked with 5% fatty acid-free bovine serum albumin (GenDEPOT, A0100) in 1% TBST (Fisher Scientific, BP2471-1, BP337-100) overnight. All the primary and secondary antibodies were diluted with 5% BSA in 1% TBST. Washes were done as a short rinse followed by 5 long washes for 5 min each. Both primary (Cell Signaling Technology, 9101, 9102) and secondary antibodies (92632211, LI-COR) were incubated for 1 h at RT. The blot was imaged with Odyssey CLx (LI-COR). The blot was stripped with Restore PLUS Western Blot Stripping Buffer (Thermo Scientific, 46430) for 15 min at RT, rinsed with 1% TBST and reblocked with 5% BSA for 30 min.

### Human Induced Pluripotent Stem Cell Derivation

Protocols were approved by the Northwestern University Institutional Review Boards. With informed written consent, ~9 ml of peripheral blood was taken from each volunteer and stored at 4 ºC, samples were transferred to Leucosep tubes (Greiner, 163288) filled with Histopaque-1077 (Sigma, 10771-100ML). 1 × 10^6^ isolated peripheral blood mononuclear cells (PMBC) were grown in 24-well tissue culture-treated plates (Greiner, 6621160) in 2 ml of SFEM II (Stem Cell Technologies, 09655) supplemented with 10 ng ml^−1^ IL3 (Peprotech, 200-03), 50 ng ml^−1^ SCF (Preprotech, 300-07), 40 ng ml^−1^ IGF1 (Peprotech, 100-11), 2 U ml^−1^ EPO (Calbiochem, 329871-50UG), 1 μM dexamethasone (Sigma, D4902) (Chou et al., 2015). 50% medium was changed every other day. After 12 days of growth, 6 × 10^4^ cells were transferred to a well of a 24-well plate in 500 μL of SFEM II with growth factors supplemented with CytoTune-iPS 2.0 Sendai Reprogramming Kit viral particle factors (Gibco, A16518) (Fujie et al., 2014; Fusaki et al., 2009) diluted to 5% (1:20) of the manufacturer’s recommendations. Cells were treated with 3.5 µL, 3.5 µl, and 2.2 µl of hKOS (0.85 × 10^8^ CIU ml^−1^), hMYC (0.85 × 10^8^ CIU ml^−1^), and hKLF4 (0.82 × 10^8^ CIU ml^−1^), respectively at MOI of 5:5:3 (KOS:MYC:KLF4). 100% media was changed after 24 h by centrifugation (300 × *g*, 4 min) to 2 ml fresh SFEM II with growth factors, and cells were transferred to one well of a 6-well plate (Greiner, 657165) coated with 1:800 Matrigel. 50% medium was changed gently every other day. On day 8 after transduction, 100% of medium was changed to B8 medium. Medium was changed every day. At day 17 individual colonies were picked into a Matrigel-coated 12-well plate (one colony per well).

### Pluripotent Cell Flow Cytometry

hiPSCs were dissociated with TrypLE Express (Gibco) for 3 min at 37 ºC and 1 × 10^6^ cells were transferred to flow cytometry tubes (Falcon, 352008). Cells were stained in 0.5% BSA in DPBS using 1:20 mouse IgG_3_ SSEA4-488 clone MC813-70 (BD Biosciences, 560308, lot 8225551) and 1:20 mouse IgM TRA-1-60-488 clone TRA-1-60 (BD Biosciences, 560173, lot 9191102) for 30 min on ice then washed. Isotype controls mouse IgG_3_-488 clone J606 (BD Biosciences, 563636, lot 7128849) and mouse IgM-488 clone G155-228 (BD Bioscience, 562409, lot 7128848) were used to establish gating. All cells were analyzed using a CytoFLEX (Beckman Coulter) with CytExpert 2.2 software.

### Immunofluorescent Staining

hiPSCs were dissociated with 0.5 mM EDTA and plated onto 1:800 Matrigel-treated Nunc Lab-Tek II 8-chamber slides in B8 medium for two days (B8T for the first 24 h). Cells were fixed with 4% PFA (Electron Microscope Sciences, 15713S) in DPBS for 15 min at RT, permeabilized with 0.3% Triton X-100 (Fisher Bioreagents, BP151-100) in DPBS for 10 min at RT, blocked with 1% BSA in DPBS for 60 min at RT, and stained with primary antibodies 1:400 polyclonal rabbit IgG anti-OCT4 (POU5F1) (Santa Cruz, sc-9081), 1:100 monoclonal mouse IgG_3_ anti-SSEA4 (Santa Cruz, sc-21704), 1:100 monoclonal rat IgG_2a_ anti-SOX2 (eBioscience, 14-9811-82), and 1:100 monoclonal mouse IgM anti-TRA-1-60 (Santa Cruz, sc-21705) in 1% BSA plus 0.1% Tween (Fisher BP337-100) for 2 h at RT. Cells were then washed three times with DPBS (2 min each), stained with secondary antibodies Alexa Fluor 594 goat anti-rabbit IgG (Invitrogen A11012), Alexa Fluor 488 goat anti-rabbit IgG (Invitrogen A21151), Alexa Fluor 488 goat anti-rat IgG (Invitrogen A11006), and Alexa Fluor 594 goat anti-rabbit IgG (Invitrogen A21044), in 1% BSA plus 0.1% Tween for 2 h at RT, washed three times with DPBS, and mounted with ProLong Diamond Antifade Mountant with DAPI (Invitrogen). Slides were imaged with a Ti-E inverted fluorescence microscope (Nikon Instruments) and a Zyla sCMOS camera (Andor) using NIS-Elements 4.4 Advanced software.

### Population Doubling Level Assessment

Population doubling level (PDL) was calculated according to the following formula:

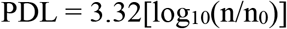

Where n = cell number and n_0_ = number of cells seeded

### Cardiac Differentiation

Differentiation into cardiomyocytes was performed according to previously described protocol with slight modifications (Burridge et al., 2015; Burridge et al., 2014). Briefly, hiPSCs were split at a 1:20 ratio using 0.5 mM EDTA as above and grown in B8 medium for 4 days reaching ~75% confluence. At the start of differentiation (day 0), B8 medium was changed to CDM3 (chemically defined medium, three components) (Burridge et al., 2014), consisting of RPMI 1640 (Corning, 10-040-CM), 500 μg ml^−1^ fatty acid-free bovine serum albumin (GenDEPOT, A0100), and 200 μg ml^−1^ L-ascorbic acid 2-phosphate (Wako, 321-44823). For the first 24 h, CDM3 medium was supplemented with 6 μM of glycogen synthase kinase 3-β inhibitor CHIR99021 (LC Labs, C-6556). On day 1, medium was changed to CDM3 and on day 2 medium was changed to CDM3 supplemented with 2 μM of the Wnt inhibitor Wnt-C59 (Biorbyt, orb181132). Medium was then changed on day 4 and every other day for CDM3. Contracting cells were noted from day 7. On day 14 of differentiation, cardiomyocytes were dissociated using DPBS for 20 min at 37 °C followed by 1:200 Liberase TH (Roche, 5401151001) diluted in DPBS for 20 min at 37 °C, centrifuged at 300 *g* for 5 min, and filtered through a 100 μm cell strainer (Falcon), and analyzed.

### Endothelial Differentiation

hiPSCs were grown to ~60% confluence and differentiated according to an adapted version of a protocol previously described (Patsch et al., 2015). On day 5 of differentiation, endothelial cells were dissociated with Accutase (Corning, 25058Cl) for 5 min at 37 °C, centrifuged at 300 *g* for 5 min, and analyzed.

### Epithelial Differentiation

hiPSCs were split at 1:20 ratio using 0.5 mM EDTA as above and grown in B8T medium for 1 day reaching ~15% confluence at the start of differentiation. Surface ectoderm differentiation was performed according to an adapted version of previously described protocols (Li et al., 2015; Qu et al., 2016). On day 4 of differentiation, epithelial cells were dissociated with Accutase for 5 min at 37 °C, centrifuged at 300 *g* for 5 min, and analyzed.

### Differentiated Cell Flow Cytometry

Cardiomyocytes were dissociated as described above and transferred to flow cytometry tubes and fixed with 4% PFA for 15 min at RT, and then permeabilized with 0.1% Triton X-100 (Fisher, BP151100) for 15 min at RT, washed once with DPBS, and stained using 1:33 mouse monoclonal IgG_1_ TNNT2-647 clone 13-11 (BD Biosciences, 565744, lot 7248637) for 30 min at RT and washed again. Isotype control mouse IgG_1_-647 clone MOPC-21 (BD Biosciences, 565571, lot 8107668) was used to establish gating. Endothelial cells were dissociated as described above, transferred to flow cytometry tubes and stained with 1:100 mouse IgG2a CD31-647 clone M89D3 (BD Bioscience, 558094, lot 8145771) for 30 min on ice then washed once with DPBS. Isotype control mouse IgG_1_-647 was used to establish gating. Epithelial cells were dissociated as described above, transferred to flow cytometry tubes, fixed with 4% PFA for 10 min at RT, and then permeabilized with 0.1% saponin in DPBS for 15 min at RT. Cells were washed once in wash buffer (DPBS with 5% FBS, 0.1% NaN_3_, 0.1% saponin), stained with 1:200 mouse IgG_1_ KRT18-647 clone DA-7 (BioLegend, 628404, lot. B208126) for 30 min at RT, then washed twice with wash buffer. Isotype control mouse IgG_1_-647 was used to establish gating. All cells were analyzed using a CytoFLEX (Beckman Coulter) with CytExpert 2.2 software.

### Statistical Methods

Data were analyzed in Excel or R and graphed in GraphPad Prism 8. Detailed statistical information is included in the corresponding figure legends. Data were presented as mean ± SEM. Comparisons were conducted via an Mann-Whitney test with significant differences defined as *P* < 0.05 (*), *P* < 0.01 (**), *P* < 0.005 (***), and *P* < 0.0001 (****). No statistical methods were used to predetermine sample size. The experiments were not randomized, and the investigators were not blinded to allocation during experiments and outcome assessment.

## Acknowledgments

This work was supported by NIH NCI grant R01 CA220002, American Heart Association Transformational Project Award 18TPA34230105, a Dixon Foundation Translational Research Grants Innovation Award (P.W.B.) and the Fondation Leducq (P.W.B., A.L.G.).

## Author Contributions

P.W.B. conceived and supervised the project. H-H.K. completed most experiments, along with K.A.F., E.A.P., C.J.W., H.F., M.V.O., M.J., M.R.-T., H.-H.K., M.B., and T.M. Recombinant protein production was completed by X.G. and plasmid generation by J.-M.D. C.E. provided technical support and A.L.G provided recombinant DNA construction. P.W.B. wrote the paper. All authors approved the manuscript.

## STAR METHODS

### KEY RESOURCES TABLE

| REAGENT or RESOURCE | SOURCE | IDENTIFIER |
| --- | --- | --- |
| **Antibodies** |  |  |
| POU5F1, polyclonal rabbit IgG | Santa Cruz | sc-9081<br>RRID:AB_2167703 |
| SSEA4, monoclonal mouse IgG<sub>3</sub> | Santa Cruz | sc-21704<br>RRID:AB_628289 |
| SOX2, monoclonal rat IgG<sub>2a</sub> | eBioscience | 14-9811-82<br>RRID:AB_11219471 |
| TRA-1-60, monoclonal mouse IgM | Santa Cruz | sc-21705<br>RRID:AB_628385 |
| Alexa Fluor 594 goat anti-rabbit IgG | Invitrogen | A11012<br>RRID:AB_141359 |
| Alexa Fluor 488 goat anti-mouse IgG<sub>3</sub> | Invitrogen | A21151<br>RRID:AB_1500907 |
| Alexa Fluor 488 goat anti-rat IgG | Invitrogen | A11006<br>RRID:AB_2534074 |
| Alexa Fluor 594 goat anti-mouse IgM | Invitrogen | A21044<br>RRID:AB_141424 |
| SSEA4-488, mouse IgG<sub>3</sub> | BD Biosciences | 560308<br>RRID:AB_1645371 |
| TRA-1-60-488, mouse IgG<sub>1</sub> | BD Biosciences | 560173<br>RRID:AB_1645379 |
| Isotype control: mouse IgG<sub>3</sub>-488 | BD Biosciences | 563636 |
| Isotype control: mouse IgM-488 | BD Biosciences | 562409 |
| TNNT2-647 | BD Biosciences | 565744<br>RRID:AB_2739341 |
| Isotype control: mouse IgG<sub>1</sub>-647 | BD Biosciences | 565571 |
| CD31-647 | BD Biosciences | 558094<br>RRID:AB_397020 |
| KRT18-647 | Biolegend | 628404<br>RRID:AB_2563650 |
| Isotype control: mouse IgG<sub>1</sub>-647 | BD Biosciences | 565571 |
| **Chemicals, Peptides, and Recombinant Proteins** |  |  |
| DMEM/F12 (with HEPES), 1 liter | Corning | 10-092-CM |
| Recombinant human insulin, | Gibco | A11382IJ |
| L -ascorbic acid 2-phosphate trisodium salt | Wako | 321-44823 |
| Recombinant human transferrin (Optiferrin) | InVitria | 777TRF029-10G |
| Sodium selenite | Sigma | S5261-10G |
| Recombinant human TGFβ1 | Peptotech | 100-21 |
| Recombinant human TGFβ3 | Cell Guidance Systems | GFH109 |
| Recombinant human NRG1 | Peptotech | 100-03 |
| Y27632 | LC Labs | Y-5301 |
| Thiazovivin | LC Labs | T-9753 |
| CHIR99021 | LC Labs | C-6556 |
| Wnt-C59 | Biorbyt | orb181132 |
| Recombinant Human Long R3 IGF1 | Sigma | 91590C |
| DPBS (without calcium and magnesium) | Corning | 21-031-CV |
| EDTA | Invitrogen | 15575020 |
| TrypLE | Gibco | 12604013 |
| Liberase TH | Roche | 5401151001 |
| Accutase | Corning | 25058CI |
| Sodium bicarbonate | Sigma | S5761 |
| NEAA | Gibco | 11140050 |
| Chemically defined lipid concentrate | Gibco | 11905031 |
| Fatty acid-free bovine albumin | GenDEPOT | A0100 |
| HCl | Sigma | H9892 |
| NaOH | Sigma | S2770 |
| Sodium chloride | Sigma | S5886 |
| Cell culture water | Corning | 25-055-CV |
| DMEM | Corning | 10-017-CV |
| Matrigel | Corning | 354230 |
| Cultrex | Trevigen | 3433-005-01 |
| One Shot BL21 Star (DE3) | Invitrogen | C601003 |
| LB Broth, Miller | Fisher | BP1425-500 |
| Kanamycin | Gibco | 11815024 |
| Ampicillin | Gibco | 11593027 |
| Magic Media | Invitrogen | K6815 |
| B-PER | Thermo | 78248 |
| HisPur Ni-NTA Spin Purification Kit | Thermo | 88229 |
| Thrombin CleanCleave Kit | Sigma | RECOMT |
| **Critical Commercial Assays** |  |  |
| MycoAlert PLUS Kit | Lonza | LT07-703 |
| PrestoBlue | Invitrogen | A13261 |
| Quant-iT Qubit Protein Assay Kit | Invitrogen | Q33211 |
| **Recombinant DNA** |  |  |
| pET28a | Novagen | 69864 |
| pET32a | Novagen | 69015 |
| FGF2-K128N | Addgene | 120284 |
| FGF2-G3 | Addgene | 120286 |
| FGF1-4X | Addgene | 120282 |
| TGFB1 | Addgene | 120361 |
| TGFB1m | Addgene | 120285 |
| TGFB3 | Addgene | 120288 |
| TGFB3m | Addgene | 120289 |
| NRG1 | Addgene | 120287 |

#### Contact for Reagent and Resource Sharing

Requests should be made to the lead contact: Paul W. Burridge, Northwestern University

Plasmids generated in this study have been deposited to Addgene (see recombinant DNA above)

---

**Figure S1.**
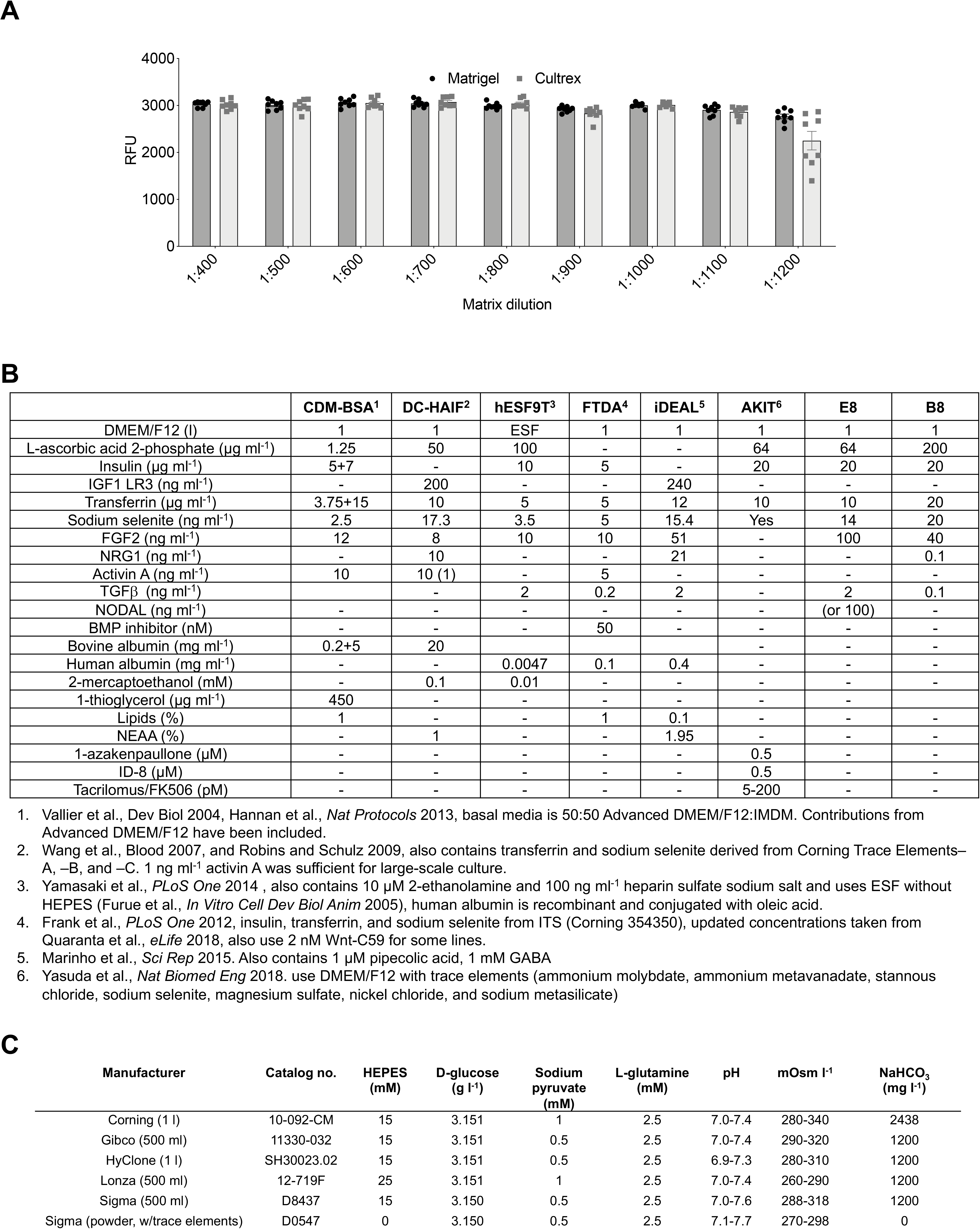
Optimization of Matrix Concentration and Comparison of Media Formulae. **a**, Relative growth of hiPSC line 19c3 on dilutions of Matrigel (Corning) or Cultrex (Trevigen), Cultrex is equivalent to Geltrex (Gibco). **b**, Comparative table of existing published media formulas for hPSC growth. **c**, Comparison of commercially available DMEM/F12 formulae that contain D-glucose, L-glutamine, phenol red, and HEPES. D0547 is the formula used by Yasuda *et al*., 2018.

**Figure S2.**
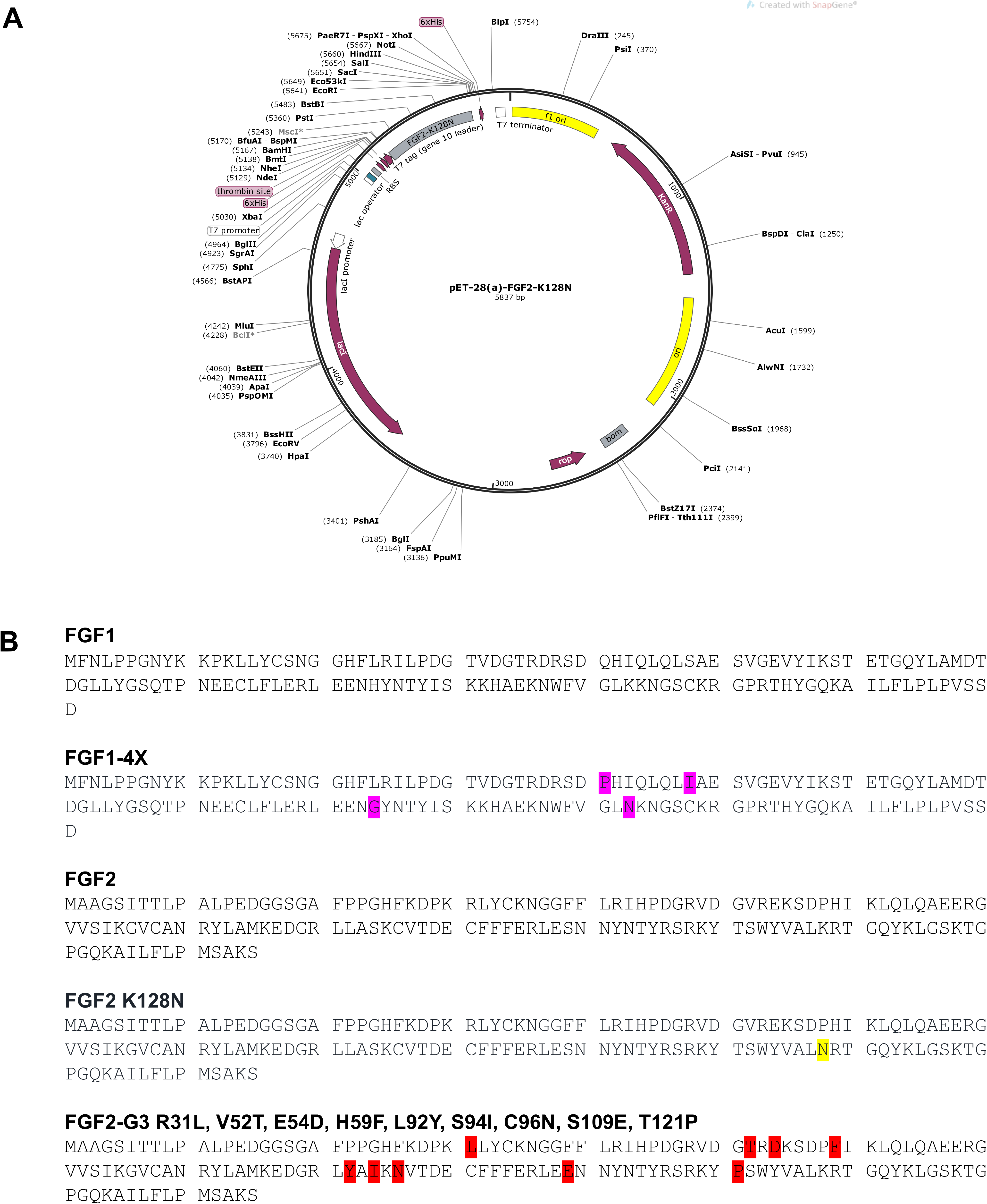
Plasmids used to Generate Recombinant Growth Factors. **(A)** FGF2-K128N demonstrating dual 6xHis site for purification and thrombin cleavage site. **(B)** Amino acid sequences used to generate modified FGF2 plasmids.

**Figure S3.**
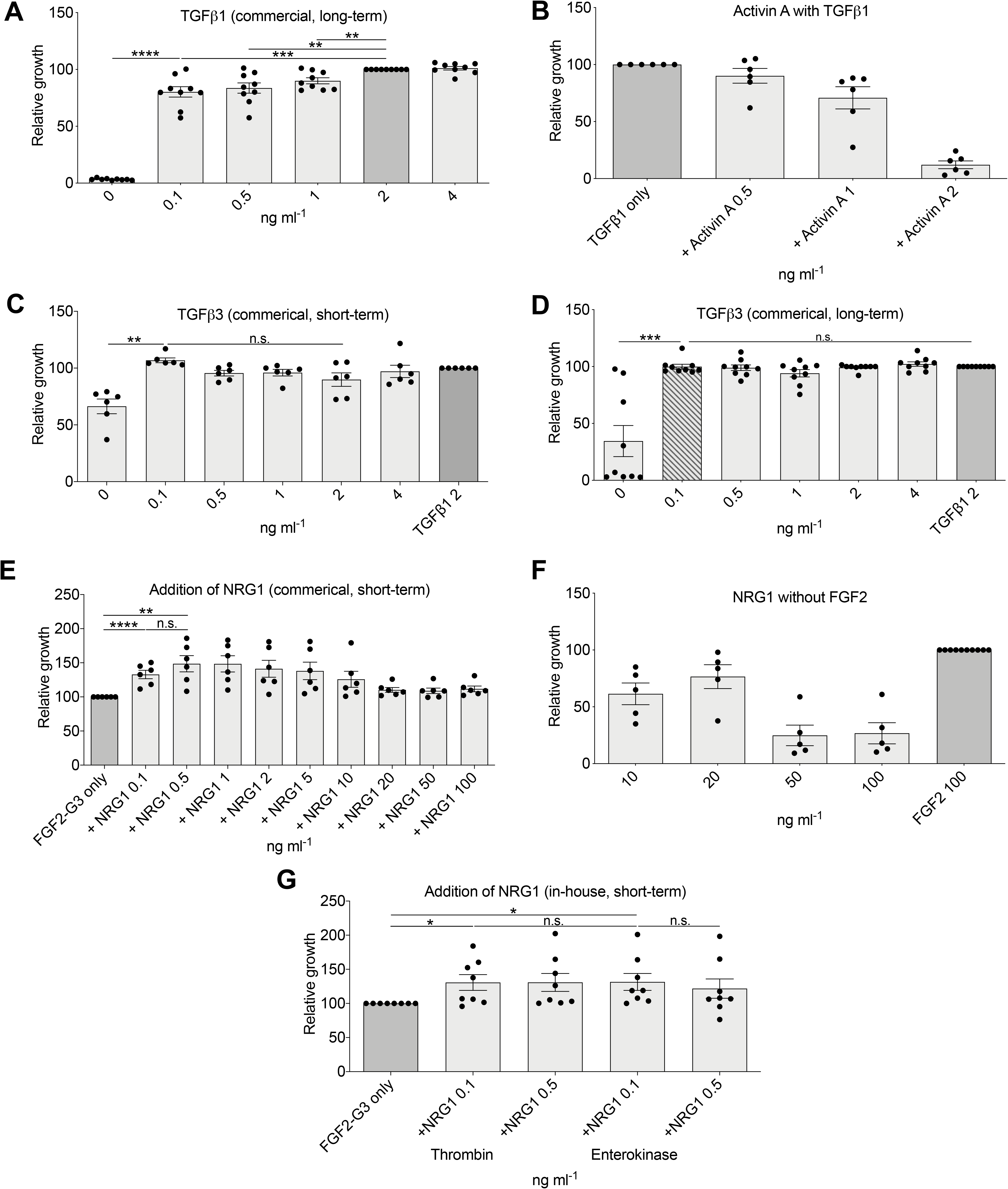
Optimization of Additional Human Pluripotent Stem Cell Medium Constituents. Results are normalized to initial medium component concentrations shown with a dark gray bar. Optimized component concentrations shown with a diagonal hash. All experiments were competed with hiPSC line 19c3 **(A)** Relative growth with commercial TGFβ1 after 9 passages (*n* = 9). **(B)** Relative growth in medium with TGFβ1 and Activin A, after 5 passages (*n* = 5). **(C)** Commercial TGFβ3 short term (*n* = 6). **(D)** Commercial TGFβ3 after 9 passages compared to commercial TGFβ1 (*n* = 9) (compare to Figure 3I, TGFβ3 (in-house). **(E)** Addition of commercial NRG1 to FGF2-G3-containing media (*n* = 6). **(F)** Relative growth in medium with NRG1 but without FGF2, long-term assay (*n* = 5). **(G)** in-house NRG1 with thrombin fusion protein cleaved by two different methods, short-term assay (*n* = 8) (compare to Figure 2J, commercial NRG1). *n* = full experimental replicates, Mann-Whitney test, \**P* ≤ 0.05, \*\**P* ≤ 0.01, \*\*\**P* ≤ 0.005, \*\*\*\**P* ≤ 0.0001, n.s. = not significant. The significance bar refers to the significance between the conditions at the beginning and the end of the bar.

**Figure S4.**
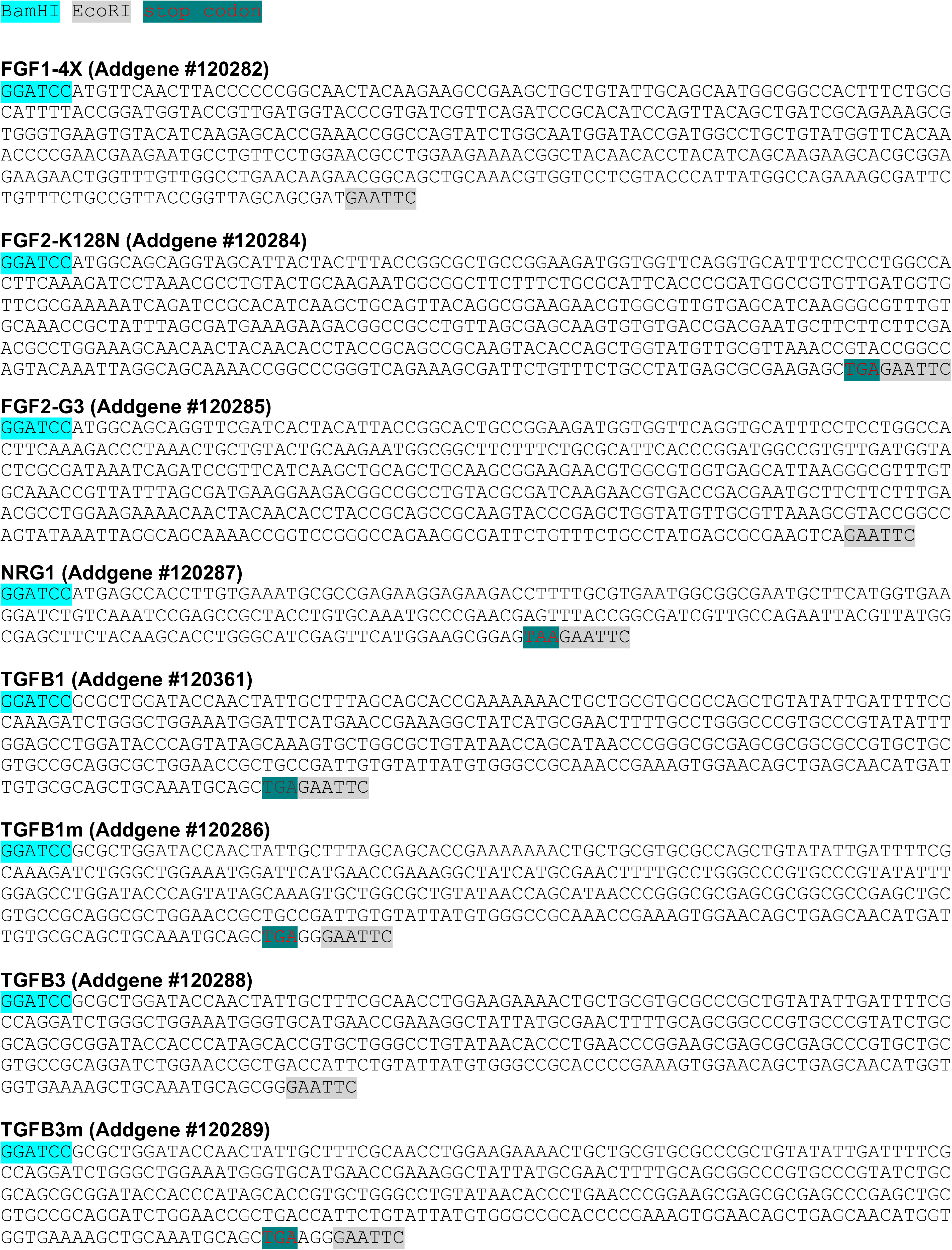
DNA Sequences used to Generate Growth Factor Plasmids. Note that FGF2-G3 and TGFβ3 were shown to function without the need for cleavage of N-terminus 6xHis tag/fusion proteins.

**Figure S5.**
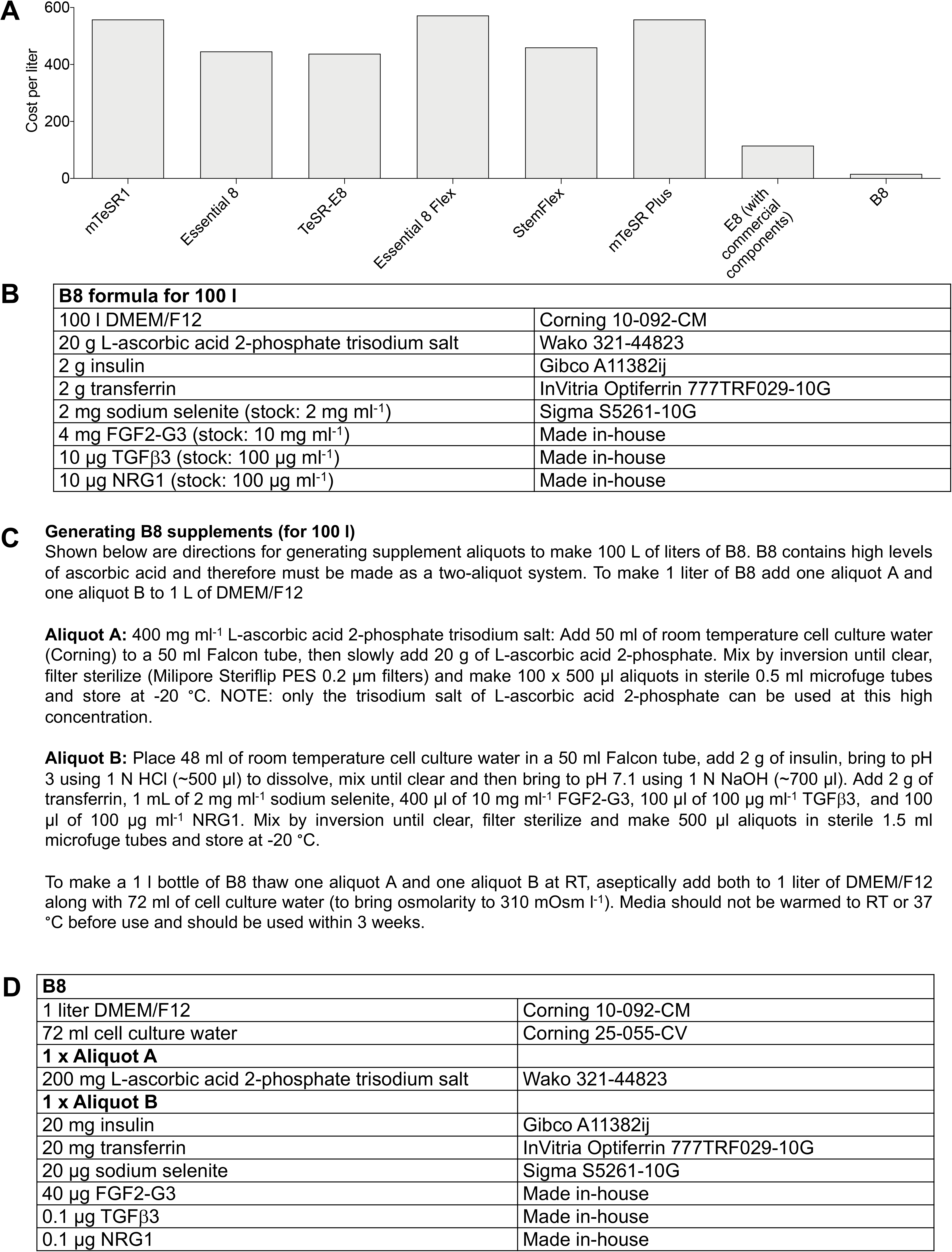
Generation of B8. **(A)** Cost comparison between B8 and commercial media. Does not include cost of shipping, technician time, or quality control. **(B)** B8 formula for 100 liters. **(C)** Short protocol for the generation of B8 supplement aliquots. **(D)** B8 formula per liter.

